# Large datasets and machine learning models fail to capture extremophile enzyme melting and optimum temperatures

**DOI:** 10.64898/2026.06.15.732300

**Authors:** Stewart Gault

## Abstract

Organisms and their enzymes adapt to environmental temperatures, such that thermophilic enzymes exhibit high melting and optimum temperatures while psychrophilic enzymes exhibit low values for both. It has been proposed that the gap between an enzyme’s optimum temperature and its melting temperature, the temperature gap, is characteristically large in psychrophiles, implying that the loss of activity above the optimum is decoupled from global protein stability. The evidence for this relies on a small number of characterised enzymes, leaving the prevalence of large temperature gaps amongst psychrophiles unknown. We asked whether the machine-learning predictors and large datasets now available could test this at scale. We find that they cannot: predictors of melting and optimum temperature fail systematically at the thermal extremes, assigning the majority of thermophilic enzymes with optimum temperatures that exceed their melting temperatures, which is biophysically implausible, and consistently underpredict the stability of (hyper)thermophiles. This stems from training data that is both error-laden, as we demonstrate for widely used optimum-temperature records, and overwhelmingly biased toward mesophiles, which regresses predictions for cold and heat adapted enzymes toward mesophilic values. Consequently, current computational tools cannot establish how prevalent the psychrophilic temperature gap is. We argue that proteome-scale measurement of extremophile enzyme thermal behaviour, integrated as curated training data, is required to determine whether trends from small studies extend across the diversity of life.

## Introduction

Over the course of Earth’s history, organisms and their enzymes have been subjected to selective pressures in adapting to new environmental temperatures and thermal regimes. As a result, organisms can be classified based on the environmental temperatures they are adapted to. Psychrophiles are adapted to grow and function at temperatures below 20 °C, with mesophiles growing between 20 to ∼ 50 °C, thermophiles grow up to 80 °C, and hyperthermophiles grow at temperatures beyond 80 °C. The enzymes of these organisms can be equally classified with their biophysical characteristics following the same trends as growth across temperature, i.e. hyperthermophilic enzymes exhibit the highest melting temperatures (*T*_m_) and optimum temperatures (*T*_opt_), while psychrophilic enzymes exhibit the lowest[1].

It has also been noted that the temperature difference between an enzyme’s *T*_opt_ and its *T*_m_, previously termed the temperature gap (*T*_gap_), is noticeably larger for psychrophilic enzymes[2], and has been suggested to be one of their defining characteristic[3]. The biophysical origin of this *T*_gap_ is still debated. Proponents of macromolecular rate theory (MMRT) argue that it is caused by the greater curvature of psychrophilic enzymes’ thermal performance curves due to their more negative Δ*C*_p_^‡^[4,5], while others argue that for psychrophilic enzymes the active site/ transition state complex is significantly more thermolabile than the bulk enzyme when compared to mesophilic or thermophilic enzymes[6–8].

Independent evidence for larger psychrophilic *T*_gap_ values can be derived from differential scanning calorimetry and neutron scattering studies[9,10]. Here, the psychrophile *Psychrobacter arcticus*’ temperature of cell death did not correlate with either the onset of thermal unfolding or the proteomes’ dynamical arrest, whereas these events were tightly coupled for the mesophile *Escherichia coli* and the hyperthermophile *Aquifex aeolicus*.

However, these studies have relied on inference from a relatively small numbers of enzymes (∼90) and organisms compared to the vast diversity that exists in nature. Here, we attempted to find evidence for the psychrophilic *T*_gap_ in large datasets and computational predictive tools of *T*_opt_ and *T*_m_ to significantly scale our understanding of psychrophilic enzymes. To our surprise we found no evidence for large psychrophilic *T*_gap_ values from predictive models and that this stems from the failure of these models to accurately capture what we consider to be “biological reality” of extremophile *T*_m_ and *T*_opt_ values. We argue that this results from poor training data that also vastly undersamples life in extreme temperature environments. Additionally, we advocate for the dedicated integration of curated and accurate extremophile protein/ enzyme biophysical and biochemical features into these large datasets.

## Results and Discussion

### The Meltome Atlas, BRENDA, and TOMER

To understand the effectiveness of large datasets and computational tools in predicting biophysical features of extremophile proteins/ enzymes it is first necessary to understand the datasets from which they are built. Thus, we started by considering the datasets of protein thermal stability.

Within the scientific literature there are many and disparate reports of protein *T*_m_ values and thermodynamics with some attempts to aggregate these into larger databases[11,12]. However, nearly every computational predictor of protein thermal stability is trained on the Meltome Atlas[13], a mass-spectrometry based *T*_m_ measurement of the entire proteome of 13 species (48,000 proteins), including one psychrophile (*Oleispira antarctica*) and three thermophiles (*Thermus thermophilus*, *Picrophilus torridus*, and *Geobacillus stearothermophilus*). The Meltome Atlas clearly shows expected features of protein biophysics; all proteins melted above the optimal growth temperature (OGT) of the respective organism, and that thermophiles have distinctly more thermostable proteomes which become more thermostable with increasing OGT. As the Meltome Atlas only contains *T*_m_ measurements it was not possible to directly measure *T*_gap_ as *T*_m_ – *T*_opt_, however, here in Figure 1 we use the OGT as a proxy for *T*_opt_. Figure 1 shows the *T*_gap_ of each species as a function of *T*_m_ – OGT, with (A) using the lowest *T*_m_ of an organism, and (B) using the median *T*_m_ value for an organism’s proteome.

**Figure 1.**
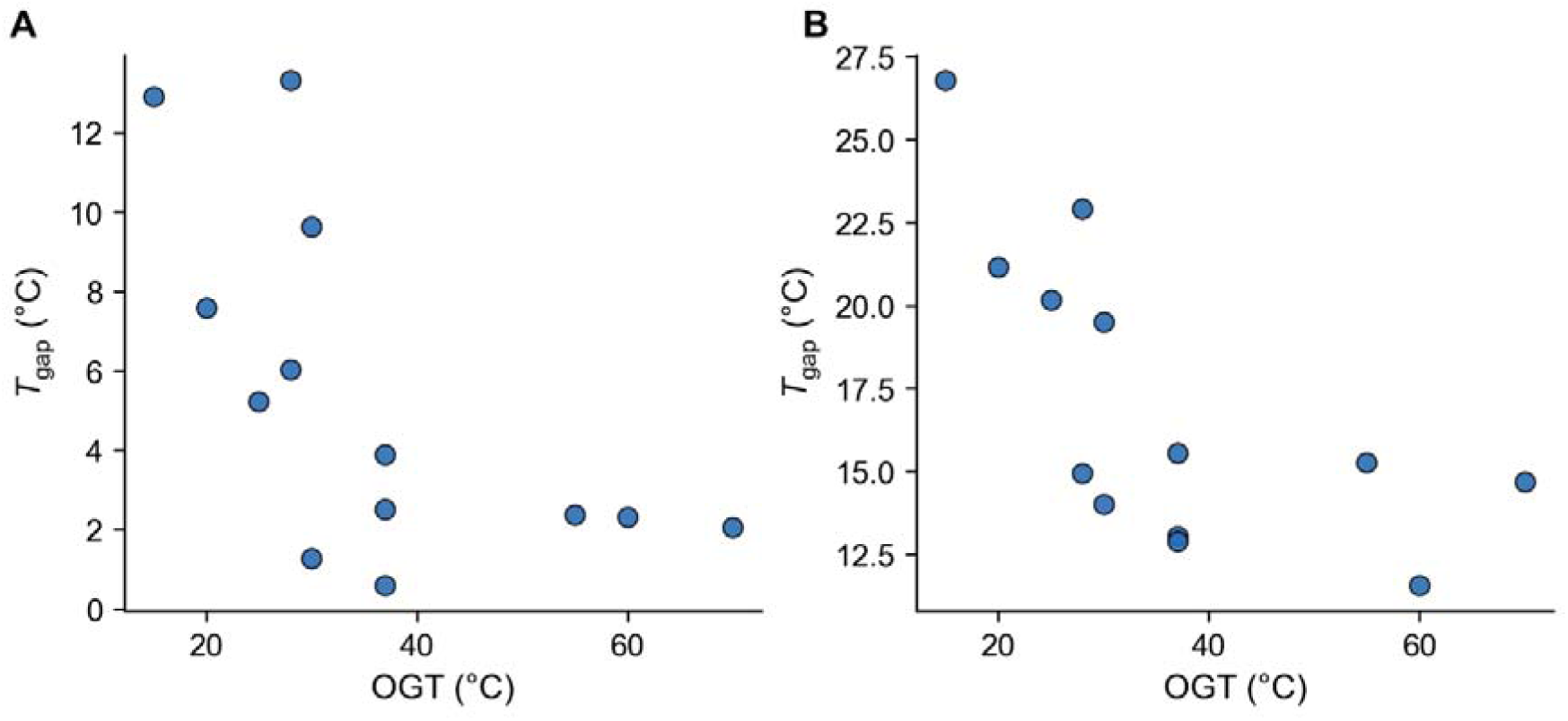
Temperature gap between *T*_m_ and an organism’s OGT. (A) shows the temperature gap between an organisms’ OGT and its lowest value for *T*_m_ i.e. the first protein to melt. (B) shows the temperature gap between the median value of an organism’s proteome *T*_m_ vs its OGT.

From Figure 1A we can see that when OGT > 35 °C the *T*_gap_ remains relatively flat, and that these species have a < 5 °C thermal buffer between their OGT and the initial onset of their proteome melt, and ∼15 °C between their OGT and the median *T*_m_ (Figure 1B). Below 35 °C however there is a clear inflection point whereupon the *T*_gap_ of a species begins to increase with decreasing OGT. Notably this inflection occurs before we reach the OGT typically ascribed to psychrophiles. However, caution is required when using OGT as a proxy for *T*_opt_. If there is effectively a minimum *T*_m_ for globular proteins at ∼ 30 ± 5°C[14–16], a floor to protein thermostability, then obligate psychrophiles with low OGT values will always present as having greater *T*_gap_ values compared to mesophiles and thermophiles. In other words, it is entirely possible for OGT to decouple from *T*_m_ without *T*_opt_ also decoupling.

Therefore, the Meltome Atlas offers tentative proteome scale evidence that organisms with low OGT values have decoupled their growth temperature optima from their proteomes’ thermostability, but the question of enzyme *T*_opt_ and *T*_gap_ remains unresolvable from the Meltome Atlas alone. We note that Figure 1A suggests that mesophiles and thermophiles have little buffering capacity between OGT and initial proteome melting, i.e. there is only marginal stability at OGT. Figure 1B demonstrates that there is a much greater capacity for thermal buffering in the majority of proteins within an organism’s proteome. This buffering capacity affects an organism’s ability to tolerate mutations that negatively affect a protein’s *T*_m_, i.e. more thermostable proteins can tolerate more substitutions[17]. This point is caveated by the fact that few organisms actually live at their OGT, with their environmental temperature (*T*_env_) typically several °C lower than their OGT (except in the case of homeotherms), thus buffering capacity is dictated by *T*_env_ and not OGT.

Our initial investigation of psychrophile *T*_gap_ values focused solely on enzymes due to the mechanistic link between the Δ*C*_p_^‡^[4,5]/ disruption of active site/ transition state architecture[6–8] inducing the loss of activity at temperatures beyond *T*_opt_ followed by the global unfolding of the protein at *T*_m_. From the Meltome Atlas it is apparent that using OGT in lieu of *T*_opt_ values produces a similar trend of larger *T*_gap_ values for cold adapted organisms. This is likely due to the behaviour of thermal performance curves across biological scales[18], where it is noted that thermal performance curves of growth rates and enzyme activity collapse to the same universal thermal performance curve. However, to determine whether the enzymes contained within the Meltome Atlas also exhibited the same *T*_gap_ behaviour, we filtered the Meltome Atlas for enzymes that could be linked to a recorded *T*_opt_ value. Enzymes were linked from the Meltome Atlas by their protein accession number to an enzyme commission number via Uniprot[19], which allowed us to parse the BRENDA database[20] for enzymes that were contained within the Meltome Atlas, but also had *T*_opt_ values recorded in BRENDA. This approach unfortunately reduces the sampling of the temperature spectrum, as neither *O*. *antarctica* nor *P. torridus* are represented in BRENDA.

Figure 2A simply shows the *T*_m_ values of the retained species across OGT, while Figure 2B shows the recorded *T*_opt_ values. It is immediately apparent from Figure 2B that many of the reported *T*_opt_ values from BRENDA are likely to be erroneous. Figure 2C shows the *T*_gap_ values for the retained species defined as *T*_m_ - *T*_opt_ across OGT. As can be seen from Figure 2C there is largely no difference in the distribution of *T*_gap_ values across OGT. Additionally, we see that negative *T*_gap_ values are recorded, which are taken to be mostly artefacts. A negative *T*_gap_ would require an enzyme to be optimally active after it has melted/ denatured which is unlikely to reflect biological reality and is more readily explained by variations in buffer composition and pH between methodologies. Alternatively, assays may genuinely report scenarios where *T*_opt_ > *T*_m_ if the unfolding of an enzyme is kinetically inhibited or if the presence of substrate increased an enzyme’s thermostability, thus allowing for the observation of *T*_opt_ > *T*_m_ over experimental timescales. In Figure 2C, small negative *T*_gap_ values likely stem from methodological/ buffer variations between measurements of *T*_opt_ and *T*_m_. Large negative *T*_gap_ values in Figure 2C are readily explained by erroneous data entries within the BRENDA database. *B. subtilis* has a *T*_gap_ of −55 °C for a pyrimidine phosphorylase that has a recorded *T*_opt_ of 99 °C which has been mined incorrectly from the primary literature[21]. Figure 2C also suggests that *T. thermophilus* exhibits the largest *T*_gap_ values, the opposite of the hypothesis that large *T*_gap_ values are diagnostic of psychrophilic enzymes. We find however that the large *T*_gap_ values for *T. thermophilus* are also driven by errors within the BRENDA database, or complications in our data extraction. For example, the *T*_gap_ of *T. thermophilus*’ phosphoglycerate kinase at 72.8 °C is driven by a recorded *T*_opt_ value of 22 °C, which in fact reflects the temperature at which the assay was conducted at, not an actual *T*_opt_[22,23]. Figure 2D shows the *T*_gap_ values across *T*_opt_, where the distinct second linear trend belongs exclusively to *T. thermophilus*, thus demonstrating the extent of its errors. Such observations expose the dangers in relying on the BRENDA database to accurately report the biochemical and biophysical properties of enzymes, something which almost every computational tool predicting such features relies on. It is also trivial to find errors for species beyond *T. thermophilus*, for example, the carbonic anhydrase from *C. elegans* is reported in BRENDA to have a *T*_opt_ of 4 °C, when this is in fact either the assay storage temperature or the centrifugation temperature (both of which are the only mention of 4 °C in the original manuscript)[24]. This is not to say that *T*_opt_ values cannot be lesser than an organism’s OGT, in fact the two are likely to be tightly correlated, but as can be seen from Figure 2B the deviations which result from errors in the BRENDA database are striking and affect further interpretation.

**Figure 2:**
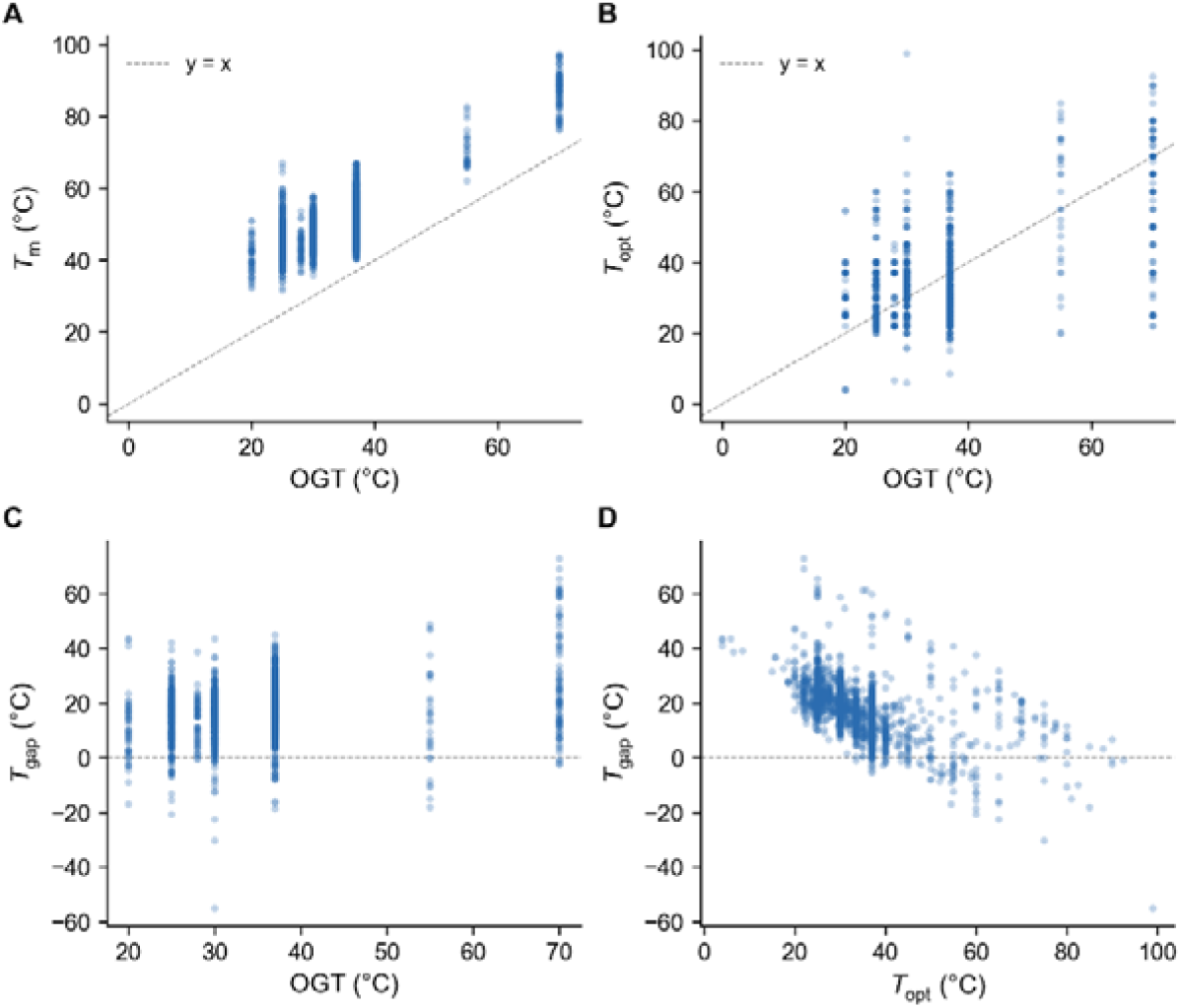
Meltome Atlas *T*_m_ and BRENDA derived *T*_opt_ values: (A) Meltome Atlas *T*_m_ values across OGT. (B) BRENDA derived *T*_opt_ values for enzymes from the Meltome Atlas. (C) shows the *T*_gap_ as *T*_m_ – *T*_opt_ across OGT with (D) showing how *T*_gap_ varies across *T*_opt_.

Attempting to link the Meltome Atlas and BRENDA to find evidence for large psychrophilic *T*_gap_ values was therefore limited on multiple fronts. Firstly, the Meltome Atlas is unrepresentative of psychrophiles with only *O. antarctica* measured, and even it had to be discounted due to its absence from the BRENDA database. Secondly, the occurrence of significant errors in BRENDA and the variety of ways in which *T*_opt_ is reported in BRENDA means simply parsing the database offers no guarantee of accuracy, meanwhile to manually curate entries severely limits the ability to scale any such analyses. However, as the Meltome Atlas and BRENDA serve as the primary sources of training data for computational predictive tools of enzyme thermal stability and activity, we wondered whether the computational tools themselves could reflect “biological reality” given the limitations of their training sets.

Currently there are a limited number of open-source tools for predicting an enzyme’s *T*_opt_, these include TOMER[25], Seq2Topt[26], and DeepET[27]. Here we chose TOMER for *T*_opt_ predictions as it requires an organism’s OGT in addition to the protein sequence, for which we used the OGT values stated in the Meltome Atlas. From Figure 3A we can see that TOMER effectively predicts two populations of *T*_opt_ values for the Meltome Atlas enzymes. For all non-thermophilic species, the median *T*_opt_ is ∼ 35 °C whereas for the thermophiles the median *T*_opt_ values are: *G. stearothermophilus* = 67 °C, *P. torridus* = 79 °C, and *T. thermophilus* = 77 °C. Additionally, TOMER did not predict any *T*_opt_ values below 29 °C. So, despite *O. antarctica* being genuinely psychrophilic with an OGT of 15 °C, TOMER does not predict that any of the computed enzymes have a similarly low *T*_opt_. We can be reasonably confident that this is a computational artefact, as it is a known issue in *T*_opt_ predictors that they overpredict psychrophilic *T*_opt_ values[26,27]. It has also been argued that much of TOMER’s predictive power stems from the inputted OGT values[26], whereas here with respect to the Meltome Atlas species it seems more likely that TOMER simply predicts mesophilic vs thermophilic traits. Figure 3B shows the resulting *T*_gap_ values from subtracting the TOMER *T*_opt_ predictions from the Meltome Atlas *T*_m_ values. It shows the same story as Figure 2C, in that when using predicted *T*_opt_ values there is essentially no correlation between the *T*_gap_ of an enzyme and its organism’s OGT. How then do we explain why experimental and computational studies have demonstrated a decoupling between a psychrophilic enzyme’s *T*_opt_ and *T*_m_ and yet predictive software does not? We believe that the answer lies directly in the quality of the training data. First consider *T*_m_, for which all predictors of *T*_m_ and thermostability are trained on a combination of both the Meltome Atlas and datasets such as FireProtDB. Both datasets are vastly unrepresentative of psychrophilic protein biophysics and as such can never be expected to truly capture the behaviour of psychrophilic proteins. Furthermore, these datasets are predominantly populated by mesophilic species and proteins which skew the psychrophilic and thermophilic predictions towards mesophily. As for *T*_opt_, the limited number of tools which are available all rely on BRENDA for training their software. However, as we have seen previously BRENDA suffers on two fronts, firstly it suffers from errors in literature extraction, or simply reports temperatures at which assays were performed at and not genuine *T*_opt_ values. Secondly it is significantly underpopulated with psychrophilic data. *T. thermophilus* alone has over 300 entries, whereas it is a challenge to find any model psychrophile with more than a handful of entries. Therefore, with the current state-of-the-art it is unlikely that scaling any of these predictors will offer much benefit to extremophile researchers.

**Figure 3:**
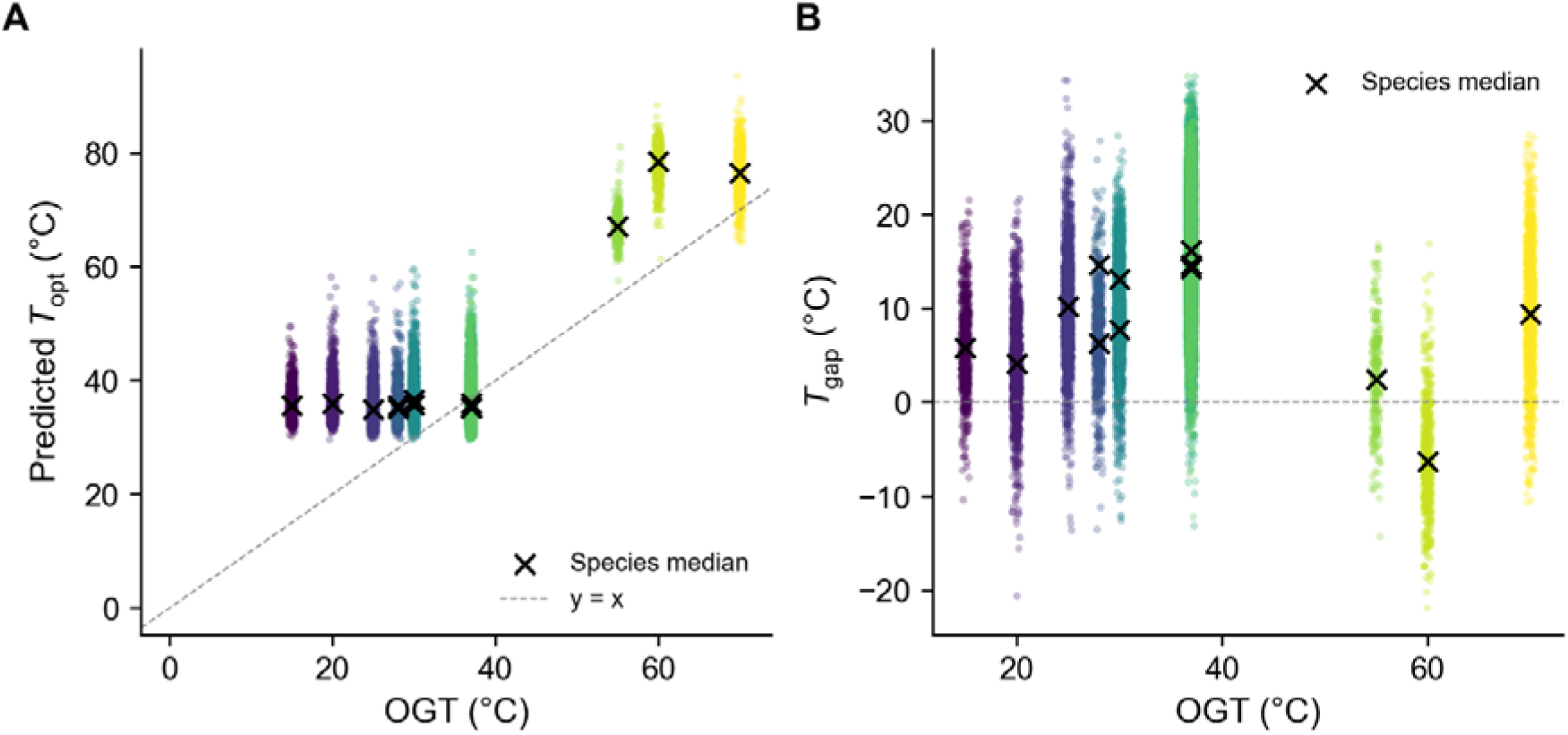
TOMER predicted *T*_opt_ values: (A) presents the TOMER predicted *T*_opt_ values across OGT while (B) shows the *T*_gap_ defined as the Meltome Atlas *T*_m_ – TOMER predicted *T*_opt_ across OGT. Each species median is highlighted with an X.

### GotEnzymes: Predictions at Scale

To explore the limitations of scaling enzyme predictions, we utilized the GotEnzymes database, consisting of computationally predicted *T*_m_, *T*_opt_, *k*_cat_, *K*_M_, and *k*_cat_/*K*_M_ values for millions of enzymes[28]. The GotEnzymes database was created by employing and improving multiple models for each of the characteristics listed previously. However, improvements in a model’s ability to predict values only reflect true improvements if the underlying data is correct.

Figure 4A shows the *T*_opt_ and *T*_m_ values for each of the enzymes contained within the GotEnzymes database, while Figure 4B displays the same but for enzymes from organisms with a reported OGT (sourced from the Gosha dataset[29]). As can be seen from Figure 4, the models employed within GotEnzymes are capable of broadly capturing the biological reality of enzyme biophysical characteristics across temperature at large scales, i.e. *T*_m_ increases with *T*_opt_. However, on closer inspection many issues are found. In the complete GotEnzymes database there are 279,369 entries where *T*_opt_ is greater than *T*_m_, representing 3.34% of database. Given that the three-dimensional structure of an enzyme is a prerequisite for activity, an enzyme exhibiting a *T*_opt_ greater than its *T*_m_ is unlikely to reflect biological reality, as previously discussed with regards to BRENDA derived values. While these values represent a minority across the entire temperature spectrum of the dataset, they become increasingly implausible at temperatures relevant to thermophiles and hyperthermophiles. We find that 74.5% of enzymes with *T*_opt_ values greater than 80 °C are predicted to have lower *T*_m_ values than their corresponding *T*_opt_ values. In other words, only ∼25% of thermophilic enzyme predictions match what we would consider to be biologically expected. As previously stated, this is an established and well-documented issue in the prediction of protein/ enzyme thermostability, arising from the overwhelming mesophile bias in training datasets which results in a significant regression to the mean where models underpredict thermophilic melting temperatures while concomitantly overpredicting the thermal stability of thermolabile (psychrophilic) proteins/ enzymes[30]. This can be seen from the fact that removing mesophilic/ low *T*_m_ proteins from datasets can improve the predictions of thermophilic *T*_m_ values[31]. The ability of these models to capture the general trends of enzyme thermostability are likely sufficient for biotechnology and de novo enzyme design purposes, but their failure to capture biological reality means they cannot be relied upon to infer how life actually behaves in extreme temperature environments.

**Figure 4:**
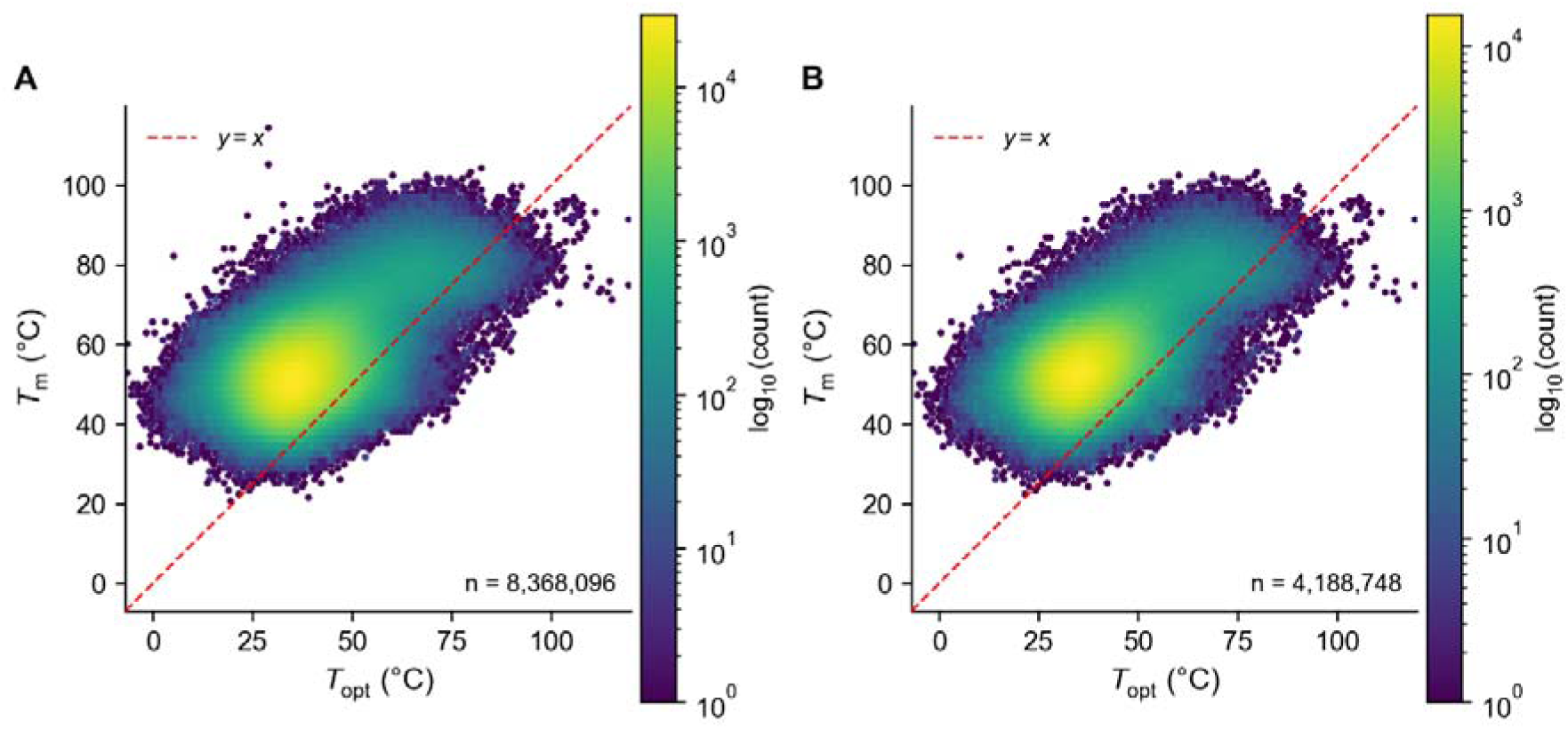
The *T*_m_ and *T*_opt_ values within the GotEnzymes database. (A) shows the *T*_m_ and *T*_opt_ values of the entire GotEnzymes database while (B) shows the same values but only for enzymes from an organism with a recorded OGT in the Gosha database. The densities of the distributions are represented on a log scale heat map.

When we try to find evidence for a larger *T*_gap_ between a psychrophile’s *T*_opt_ and its *T*_m_, the GotEnzymes dataset shows no correlation between the OGT of an organism and its enzymes’ *T*_gap_ values as demonstrated in Figure 5A. This is not unexpected from the GotEnzymes data as *T*_m_ and *T*_opt_ both exhibit linear relationships with regards to OGT as shown in Figure 5C and D respectively, and as such it is impossible for them to have ever captured a non-linear relationship. This is equally reflected in Figure 5B, where, like with the Meltome Atlas, we replace *T*_opt_ with OGT to determine *T*_gap_ which reveals a negative correlation of *T*_gap_ with respect to OGT. Across the whole temperature range this results in 1.26 % of entries having enzymes whose organisms have OGT values greater than their enzymes’ *T*_m_ values. When we solely consider the hyperthermophiles, i.e. organisms with OGT > 80 °C, this percentage increases to 79.62 %. This is simply a consequence of models underpredicting *T*_m_ values from thermostable enzymes. Indeed, if we instead consider the median *T*_m_ and *T*_opt_ values per organism, as seen in Supplementary Figure 1, the picture is even more damning. When collapsed to median per organism, 94.2 % of species with an OGT > 80 °C have median *T*_m_ values lower than their OGT while 100 % of them have *T*_opt_ values lesser than their OGT. As can be seen from Supplementary Figure 1 the departure from *T*_x_ = OGT gets progressively worse as OGT increases, showing that the predictive software fundamentally cannot capture hyperthermophile biophysics. We can be confident that this is a departure from biological reality, as it is unlikely that hyperthermophiles would contain enzymes that melt/ denature at temperatures below their OGT. This is reflected in the Meltome Atlas data analysed in Figure 2A, where the first protein to melt always exceeded the OGT of the respective organism and the median *T*_m_ was greater again.

**Figure 5.**
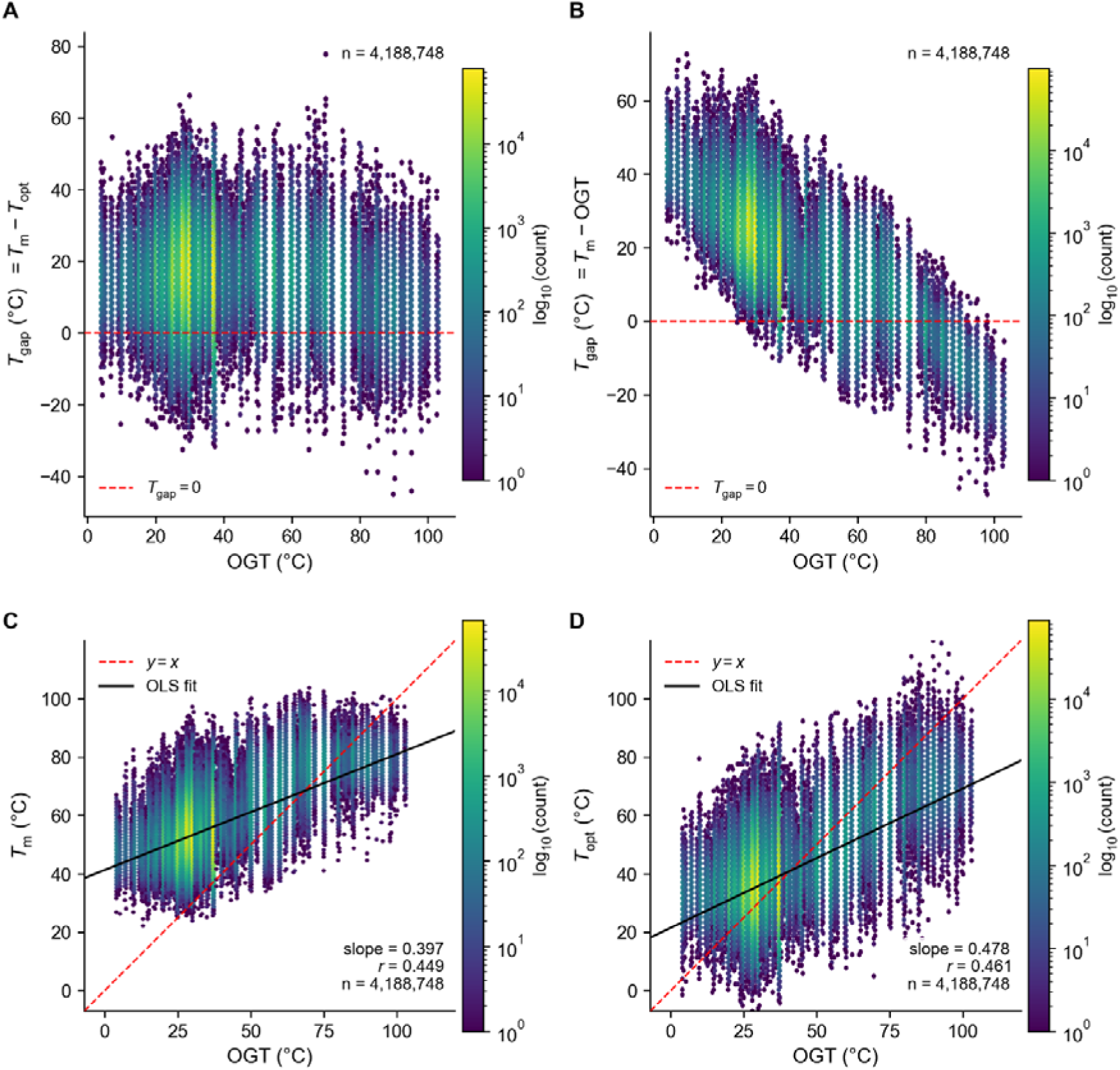
*T*_gap_ values predicted from the GotEnzymes database. (A) shows the *T*_gap_ values defined as *T*_m_ – *T*_opt_ predicted from the GotEnzymes database across OGT while (B) shows *T*_gap_ defined as *T*_m_ – OGT across OGT. (C) shows the GotEnzymes *T*_m_ values across OGT and (D) presents *T*_opt_ values across OGT. The densities of the distributions are represented on a log scale heat map.

### The Extreme Environment Microbiome Catalog

One of the motivations for pursuing this study was to determine whether computational predictors of *T*_opt_ and *T*_m_ can confidently be applied to large metagenomic datasets, in particular the Extreme Environment Microbiome Catalog (EEMC)[32]. The EEMC comprises of reconstructed microbial genomes from seven classes of extreme environments: cryosphere, deep-sea, acidic, arid, hypersaline, subsurface, and terrestrial geothermal. If such tools could be reliably applied to environmental scale databases, then it would have been possible to test the hypothesis that psychrophiles have larger *T*_gap_ values by expecting to see larger *T*_gap_ values in cryosphere and deep-sea environments. Figure 6A and B shows how organisms from the seven extreme environments are separated by amino acid Z scores, with Figure 6C showing the predicted *T*_m_ values of 100,000 proteins from each environment, predicted via TmProt[33] as it does not require OGT as an input. In Figure 6A and 6B the EEMC environments are plotted against Z scores of two different amino acid compositions; EIKPRVY[34], and IVYWREL[35], both of which have been proposed to correlate with thermal adaptation. We can see that the cryosphere exhibits the lowest Z score across both compositions, with terrestrial geothermal environments exhibiting the highest Z scores. However, when we compare this to the predicted *T*_m_ values from TmProt in Figure 6C we can see that nearly every environment has a bimodal distribution of *T*_m_ values. Each environment has a major density of *T*_m_ values around 50 °C, with a second distribution around 70 °C. This is again due to the nature of the training data for TmProt which utilised the Meltome Atlas and ProThermDB, which as previously discussed essentially means that these types of software can predict whether a protein is mesophilic or thermophilic, but that they fail to capture a smooth distribution. Figure 6D and E show how the organism Z scores and the organism’s mean *T*_m_ values are distributed when plotted together. Given the inaccuracy of *T*_opt_ predictors, we opted to not apply them to the EEMC data, or expand *T*_m_ measurements to the entire EEMC.

**Figure 6:**
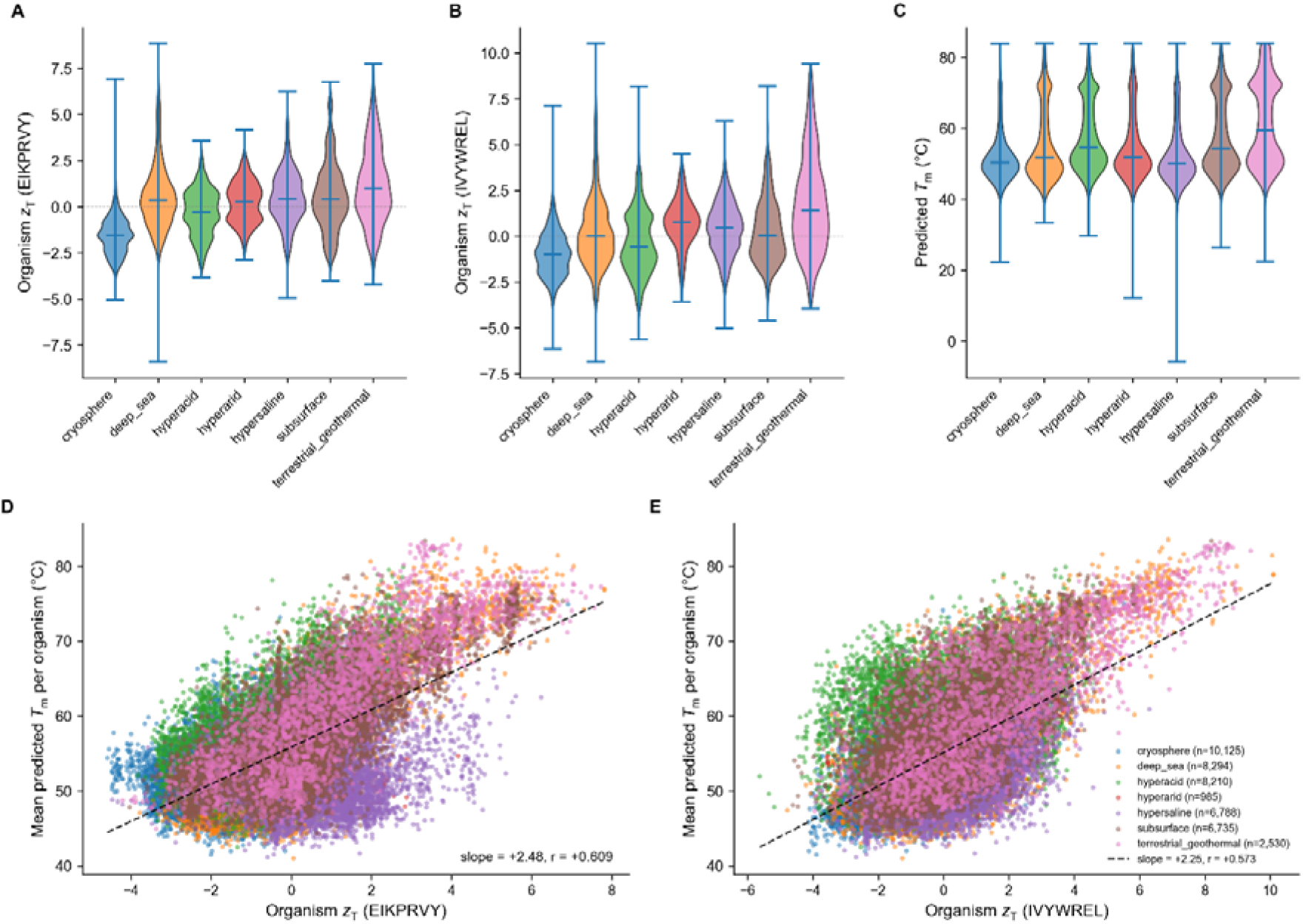
EEMC Z scores and *T*_m_ predictions. (A) shows the Z score distributions of each of the EEMC extreme environments using EIKPRVY as the amino acids, while (B) shows the distributions when IVYWREL is used. (C) shows the TmProt predicted *T*_m_ values for 100,000 randomly sampled proteins from each environment with lengths between 100-1500 amino acids. (D) and (E) show the Z scores of EIKPRVY and IVYWREL respectively plotted against the predicted *T*_m_ values.

From the data presented here it is clear that computational predictors of protein/ enzyme thermostability and activity struggle to reflect what would be considered biological reality. This is unambiguously a result of the limited training data, with the Meltome Atlas and ProThermDB/ FireProtDB datasets being unrepresentative of the thermal breadth of life. Additionally, the strong mesophilic overrepresentation biases the software against the high or low values we expect of thermophiles and psychrophiles. To remedy this, we need a concerted effort to acquire proteome-scale melting behaviour of extremophiles to greatly expand the representativeness of the Meltome Atlas. While simple in principle, it is experimentally time consuming, and care would be required in the selection of model species. Ideally a mixture of bacteria and archaea from the cryosphere and hydrothermal environments should be selected to account for amino acid biases in their proteomes across thermal scales. Additional confounders such as high pressures or high salinities would need to be accounted for before ascribing the melting behaviour and amino acid composition as being purely in response to thermal adaptation. The path forward for predictions of *T*_opt_ is slightly more nuanced. Software which rely on parsing BRENDA for temperature optima values would benefit from excluding any values which have comments such as “assayed at” to avoid erroneous *T*_opt_ entries or utilise agentic models to check source data at scale to confirm whether *T*_opt_ values were correctly extracted. Experimentally, methodologies such as high-throughput microfluidic enzyme kinetics (HT-MEK)[36] may allow for the rapid exploration of enzyme activity across temperature and assay conditions which would no doubt provide a wealth of biochemical and biophysical data for extremophile researchers.

With regards to the *T*_gap_ hypothesis that motivated this study, it seems that for now we are still reliant on experimental data for understanding the extent to which psychrophilic organisms and their enzymes are decoupled from the global stability of their proteins. As can be seen from the Meltome Atlas and previous studies[37], if the minimum temperature for protein unfolding is ∼30 ± 5 °C, then relying on OGT or *T*_env_ will always result in psychrophiles, particularly obligate psychrophiles, appearing to have a larger *T*_gap_. However this does not mean that their enzymes’ *T*_opt_ values are equally decoupled from *T*_m_ as even though the thermal performance curves converge to the same shape across scales[18], the decline in growth rates above OGT could just as well be ascribed to a hierarchical scale between enzyme activity and microbial growth breaking down which would be better described by systems biology.

## Methods and data

The datasets used in this analysis are freely available and were obtained from their respective online sources. The Meltome Atlas[13] data was downloaded from (https://meltomeatlas.proteomics.wzw.tum.de/master_meltomeatlasapp/), and the BRENDA database[20] used was 2026.1. The TOMER [25] python package and requirements were downloaded from (https://github.com/jafetgado/tomer). The data for GotEnzymes2[28] (https://github.com/LiLabTsinghua/GotEnzymes2/tree/main) was downloaded from Zenodo (DOI: 10.5281/zenodo.17168793) with the OGT values obtained from the Gosha database[29] (DOI: <u>10.6084/m9.figshare.23257940</u>). The EEMC[32] data was accessed from (DOI: <u>10.26036/CNP0007106</u>) with the code for TmProt[33] downloaded from (https://github.com/loschmidt/TmProt).

## Data availability

All data used to produce the figures in this paper are freely available at Zenodo, DOI: 10.5281/zenodo.20614369

## Notes

https://zenodo.org/records/20614369

### Competing Interest Statement

The authors have declared no competing interest.

## References

1. Siddiqui KS, Cavicchioli R. Cold-Adapted Enzymes. Annu Rev Biochem. 2006;75: 403–433. doi:10.1146/annurev.biochem.75.103004.142723

2. D’Amico S, Marx J-C, Gerday C, Feller G. Activity-stability relationships in extremophilic enzymes. J Biol Chem. 2003;278: 7891–7896. doi:10.1074/jbc.M212508200

3. Gault S, Higgins PM, Cockell CS, Gillies K. A meta-analysis of the activity, stability, and mutational characteristics of temperature-adapted enzymes. Biosci Rep. 2021;41: BSR20210336. doi:10.1042/BSR20210336

4. Hobbs JK, Jiao W, Easter AD, Parker EJ, Schipper LA, Arcus VL. Change in heat capacity for enzyme catalysis determines temperature dependence of enzyme catalyzed rates. ACS Chem Biol. 2013;8: 2388–2393. doi:10.1021/cb4005029

5. Arcus VL, Prentice EJ, Hobbs JK, Mulholland AJ, Van der Kamp MW, Pudney CR, et al. On the Temperature Dependence of Enzyme-Catalyzed Rates. Biochemistry. 2016;55: 1681–1688. doi:10.1021/acs.biochem.5b01094

6. Åqvist J, Brandsdal BO. Computer Simulations of the Temperature Dependence of Enzyme Reactions. J Chem Theory Comput. 2025;21: 1017–1028. doi:10.1021/acs.jctc.4c01733

7. Sočan J, Purg M, Åqvist J. Computer simulations explain the anomalous temperature optimum in a cold-adapted enzyme. Nat Commun. 2020;11: 2644. doi:10.1038/s41467-020-16341-2

8. van der Ent F, Yu S, Lund BA, Brandsdal BO, Sheng X, Åqvist J. Computational Design of Highly Efficient Cold-Adapted Enzymes with Elevated Temperature Optima. ACS Catal. 2025;15: 11257–11265. doi:10.1021/acscatal.5c02643

9. Caviglia B, Timr S, Guiral M, Giudici-Orticoni M-T, Seydel T, Beck C, et al. Cytoplasmic fluidity and the cold life: proteome stability is decoupled from viability in psychrophiles. Nat Commun. 2025;16: 10345. doi:10.1038/s41467-025-65270-5

10. Caviglia B, Di Bari D, Timr S, Guiral M, Giudici-Orticoni M-T, Petrillo C, et al. Decoding the Role of the Global Proteome Dynamics for Cellular Thermal Stability. J Phys Chem Lett. 2024;15: 1435–1441. doi:10.1021/acs.jpclett.3c03351

11. Stourac J, Dubrava J, Musil M, Horackova J, Damborsky J, Mazurenko S, et al. FireProtDB: database of manually curated protein stability data. Nucleic Acids Res. 2021;49: D319–D324. doi:10.1093/nar/gkaa981

12. Nikam R, Kulandaisamy A, Harini K, Sharma D, Gromiha MM. ProThermDB: thermodynamic database for proteins and mutants revisited after 15 years. Nucleic Acids Res. 2021;49: D420–D424. doi:10.1093/nar/gkaa1035

13. Jarzab A, Kurzawa N, Hopf T, Moerch M, Zecha J, Leijten N, et al. Meltome atlas—thermal proteome stability across the tree of life. Nat Methods. 2020;17: 495–503. doi:10.1038/s41592-020-0801-4

14. Martin OA, Vila JA. The Marginal Stability of Proteins: How the Jiggling and Wiggling of Atoms is Connected to Neutral Evolution. J Mol Evol. 2020;88: 424–426. doi:10.1007/s00239-020-09940-6

15. Pastore A, Martin SR, Politou A, Kondapalli KC, Stemmler T, Temussi PA. Unbiased Cold Denaturation: Low- and High-Temperature Unfolding of Yeast Frataxin under Physiological Conditions. J Am Chem Soc. 2007;129: 5374–5375. doi:10.1021/ja0714538

16. Roman EA, Faraj SE, Cousido-Siah A, Mitschler A, Podjarny A, Santos J. Frataxin from Psychromonas ingrahamii as a model to study stability modulation within the CyaY protein family. Biochimica et Biophysica Acta (BBA) - Proteins and Proteomics. 2013;1834: 1168–1180. 10.1016/j.bbapap.2013.02.015

17. Bloom JD, Labthavikul ST, Otey CR, Arnold FH. Protein stability promotes evolvability. Proc Natl Acad Sci U S A. 2006;103: 5869–5874. doi:10.1073/pnas.0510098103

18. Arnoldi J-F, Jackson AL, Peralta-Maraver I, Payne NL. A universal thermal performance curve arises in biology and ecology. Proceedings of the National Academy of Sciences. 2025;122: e2513099122. doi:10.1073/pnas.2513099122

19. Consortium TU. UniProt: the Universal Protein Knowledgebase in 2025. Nucleic Acids Res. 2025;53: D609–D617. doi:10.1093/nar/gkae1010

20. Hauenstein J, Jeske L, Jäde A, Krull M, Dümmer K, Koblitz J, et al. BRENDA in 2026: a Global Core Biodata Resource for functional enzyme and metabolic data within the DSMZ Digital Diversity. Nucleic Acids Res. 2026;54: D527–D534. doi:10.1093/nar/gkaf1113

21. Kamel S, Thiele I, Neubauer P, Wagner A. Thermophilic nucleoside phosphorylases: Their properties, characteristics and applications. Biochimica et Biophysica Acta (BBA) - Proteins and Proteomics. 2020;1868: 140304. doi:10.1016/j.bbapap.2019.140304

22. Littlechild JA, Davies GJ, Gamblin SJ, Watson HC. Phosphoglycerate kinase from the extreme thermophile Thermus thermophilus Crystallization and preliminary X-ray data. FEBS Lett. 1987;225: 123–126. doi:10.1016/0014-5793(87)81143-2

23. Nojima H, Oshima T, Noda H. Purification and Properties of Phosphoglycerate Kinase from Thermus thermophilus Strain HB8. The Journal of Biochemistry. 1979;85: 1509–1516. doi:10.1093/oxfordjournals.jbchem.a132480

24. Fasseas MK, Tsikou D, Flemetakis E, Katinakis P. Molecular and biochemical analysis of the α class carbonic anhydrases in Caenorhabditis elegans. Mol Biol Rep. 2011;38: 1777–1785. doi:10.1007/s11033-010-0292-y

25. Gado JE, Beckham GT, Payne CM. Improving Enzyme Optimum Temperature Prediction with Resampling Strategies and Ensemble Learning. J Chem Inf Model. 2020;60: 4098–4107. doi:10.1021/acs.jcim.0c00489

26. Qiu S, Hu B, Zhao J, Xu W, Yang A. Seq2Topt: a sequence-based deep learning predictor of enzyme optimal temperature. Brief Bioinform. 2025;26: bbaf114. doi:10.1093/bib/bbaf114

27. Li G, Buric F, Zrimec J, Viknander S, Nielsen J, Zelezniak A, et al. Learning deep representations of enzyme thermal adaptation. Protein Science. 2022;31: e4480. 10.1002/pro.4480

28. Lyu B, Wu K, Huang Y, Anton M, Li X, Viknander S, et al. GotEnzymes2: expanding coverage of enzyme kinetics and thermal properties. Nucleic Acids Res. 2026;54: D583–D592. doi:10.1093/nar/gkaf1053

29. Helena-Bueno K, Brown CR, Melnikov S. Gosha: a database of organisms with defined optimal growth temperatures. bioRxiv. 2023; 2021.12.21.473645. doi:10.1101/2021.12.21.473645

30. Rodella C, Lazaridi S, Lemmin T. TemBERTure: advancing protein thermostability prediction with deep learning and attention mechanisms. Bioinformatics Advances. 2024;4: vbae103. doi:10.1093/bioadv/vbae103

31. Li M, Wang H, Yang Z, Zhang L, Zhu Y. DeepTM: A deep learning algorithm for prediction of melting temperature of thermophilic proteins directly from sequences. Comput Struct Biotechnol J. 2023;21: 5544–5560. 10.1016/j.csbj.2023.11.006

32. Jiang P, Liang Z, Kovacevic V, Shi J, Milicevic N, Wang F, et al. The Extreme Environment Microbiome Catalog (EEMC): a global resource for microbial diversity and antimicrobial discovery. Nat Commun. 2026;17: 4791. doi:10.1038/s41467-026-71145-0

33. Pailozian K, Kohout P, Damborsky J, Mazurenko S. Investigation of Protein Melting Temperature Prediction with Cross-Method Validation on Biophysical Data. bioRxiv. 2026; 2026.05.07.723192. doi:10.64898/2026.05.07.723192

34. Amangeldina A, Tan ZW, Berezovsky IN. Living in trinity of extremes: Genomic and proteomic signatures of halophilic, thermophilic, and pH adaptation. Curr Res Struct Biol. 2024;7: 100129. 10.1016/j.crstbi.2024.100129

35. Zeldovich KB, Berezovsky IN, Shakhnovich EI. Protein and DNA Sequence Determinants of Thermophilic Adaptation. PLoS Comput Biol. 2007;3: e5-. doi:10.1371/journal.pcbi.0030005

36. Markin CJ, Mokhtari DA, Sunden F, Appel MJ, Akiva E, Longwell SA, et al. Revealing enzyme functional architecture via high-throughput microfluidic enzyme kinetics. Science (1979). 2021;373: eabf8761. doi:10.1126/science.abf8761

37. Dehouck Y, Folch B, Rooman M. Revisiting the correlation between proteins’ thermoresistance and organisms’ thermophilicity. Protein Engineering, Design and Selection. 2008;21: 275–278. doi:10.1093/protein/gzn001

