## Supplementary figures and images for "Large datasets and machine learning models fail to capture extremophile enzyme melting and optimum temperatures"

### Supplementary data

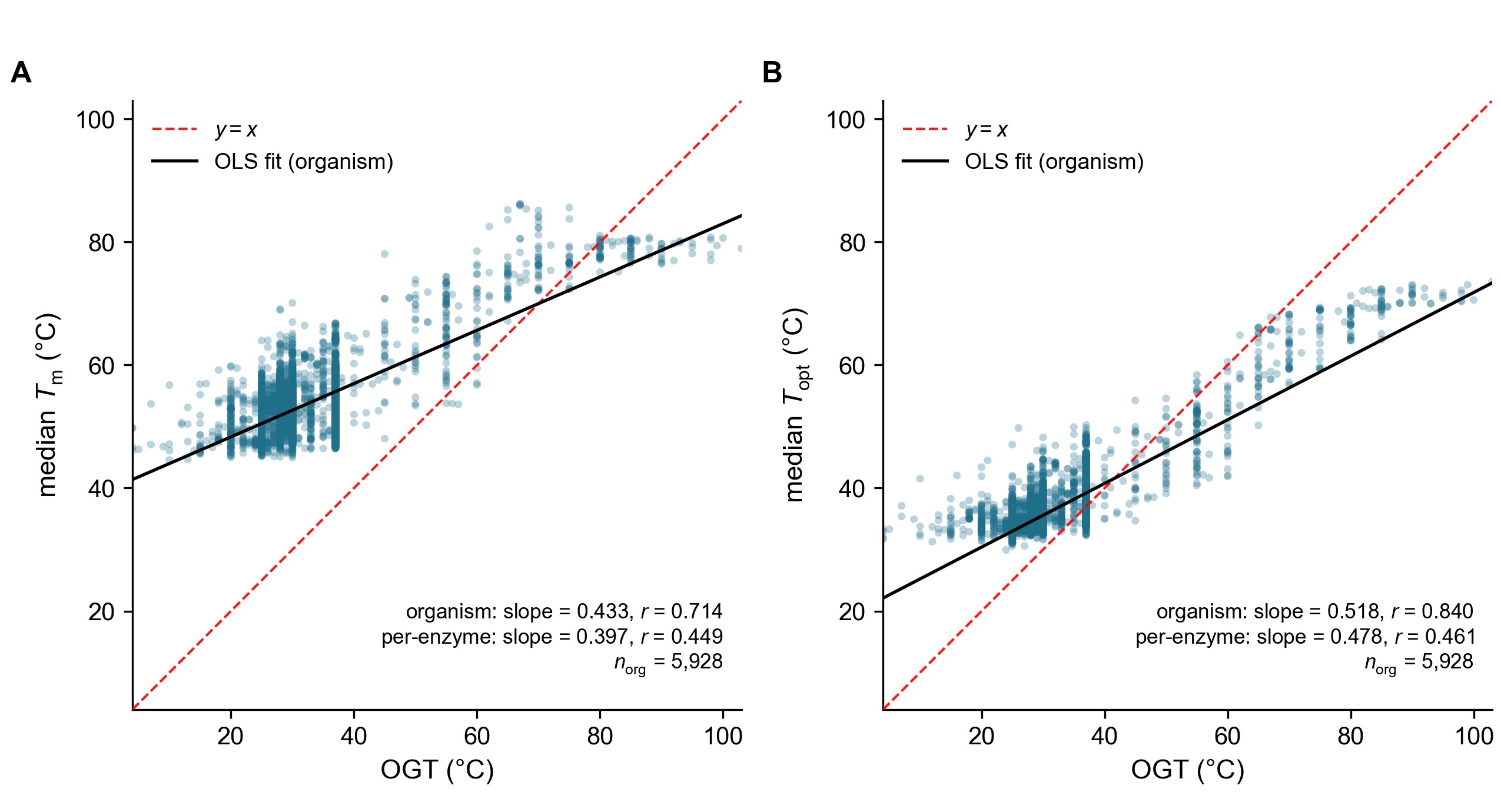
